# Dynamical modulation of theta-gamma coupling during REM sleep

**DOI:** 10.1101/169656

**Authors:** Mojtaba Bandarabadi, Richard Boyce, Carolina Gutierrez Herrera, Claudio Bassetti, Sylvain Williams, Kaspar Schindler, Antoine Adamantidis

**Affiliations:** Department of Neurology, Inselspital University Hospital, University of Bern, Bern, Switzerland; Department for BioMedical Research, University of Bern, Bern, Switzerland; Integrated Program in Neuroscience, McGill University, Montreal, Quebec, Canada; Department of Psychiatry, McGill University, Montreal, Quebec, Canada

**Author notes:** Co-last authors (Kaspar Schindler, Antoine Adamantidis). Correspondence to*: Antoine Adamantidis, Ph.D., Department of Neurology, Inselspital University Hospital, University of Bern, Freiburgstrasse 18, 3010 Bern, Switzerland.

## Abstract

Theta phase modulates gamma amplitude during spatial navigation and rapid eye movement sleep (REMs). Although the REMs theta rhythm has been linked to spatial memory consolidation, the underlying mechanism remains unclear. We investigate dynamics of theta-gamma interactions across multiple frequency and temporal scales in simultaneous recordings from hippocampal CA3, CA1, subiculum, and parietal cortex. We show that theta phase significantly modulates three distinct gamma bands during REMs, dynamically. Interestingly, we further show that theta-gamma coupling swings between different hippocampal and cortical structures during REMs and tends to increase over a single REMs episode. Comparing to active wake, theta-gamma coupling during REMs is significantly increased for subicular and cortical, but not for CA3 and CA1, recordings. Finally, optogenetic silencing of septohippocampal GABAergic projections significantly impedes both theta-gamma coupling and theta phase coherence, two neural mechanisms of working and long-term memory, respectively. Thus, we show that theta-gamma coupling and theta phase coherence are highly modulated during single REMs episode and propose that theta-gamma coupling provides a predominant mechanism for information processing within each brain region, while the orchestrated nature of coupling activity establishes a specific phase-space coding of information during sleep.

## Introduction

Both rapid eye movement sleep (REMs) and non-REMs (NREMs) are associated with consolidation of some aspects of memory following task acquisition in rodents and humans^1-5^. Classically, the reactivation of hippocampal place cells during NREMs provides one possible mechanism for the long-term encoding of newly acquired spatial^1,6^, but also procedural and emotional information^7,8^. However, the underlying mechanisms of memory consolidation during REMs is unclear. Recent work involving in-depth analysis of local field potential (LFP < 500 Hz) signals recorded from key memory-associated brain structures has revealed important insight into the neural circuitry and specific neural mechanisms that likely mediate memory consolidation^9-11^.

Indeed, phase-amplitude cross-frequency coupling, which describes the modulation by slow oscillation phase of fast oscillation amplitude, has been shown to be increased during cognitive and perceptual tasks^12,13^. Specifically, hippocampal and cortical phase-amplitude theta-gamma coupling is prevailing during locomotion and REMs in rodents^14,15^, and increases following behavioural and learning tasks in rodents^16,17^, monkeys^18,19^, and humans^20,21^. In addition, dynamical phase entrainment of low frequency oscillations (<15 Hz) during cognitive tasks and sensory inputs has also been described^22,23^. Thus, the existence of phase-amplitude coupling and phase entrainment of low frequencies demonstrates the broad functional role of phase-amplitude coupling in hierarchical information processing essential to cognitive functions (for a review^24^). Furthermore, impaired phase-amplitude coupling has been reported in Alzheimer’s disease^25,26^, schizophrenia^27^ and Parkinson’s disease^28^.

The hippocampal formation is essential to memory, learning, and fundamental behaviours such as spatial navigation^29^. Theta and gamma are predominant oscillations during active wake (aWk) and REMs in rodent hippocampal structures, and are increased by spatial navigation and learning tasks^30^. Theta rhythm in the hippocampus is generated through extrinsic^3,31^ and intrinsic mechanisms^32,33^. We recently showed for the first time direct evidence that the hippocampal theta rhythm during REMs is involved in contextual memory consolidation^3^. By suppressing extrinsic theta-rhythmic projections into the hippocampus during REMs after learning through optogenetic silencing of medial septal GABAergic (MS^GABA^) neurons, we demonstrated impaired object recognition and fear-conditioned contextual memory in mice. However, how the REMs theta rhythm contributes to memory formation is unclear.

Here, we investigated the function of REMs theta rhythm in information processing by exploring phase-amplitude coupling between theta and fast oscillations in hippocampal LFP and parietal electrocorticographic (ECoG) recordings of freely-moving mice during spontaneous sleep-wake states and upon optogenetic perturbations of MS^GABA^ neurons during REMs. We found that episodes of both aWk and REMs showed highly significant theta-middle gamma coupling, with the highest level of coupling observed during REMs. Theta-high gamma and theta-ultrahigh gamma coupling was also present during REMs, but not during aWk. We further found that theta-gamma coupling is a discrete phenomenon that swings between hippocampal/cortical structures, and increases over REMs. Finally, optogenetic silencing of MS^GABA^ neurons impeded phase-amplitude coupling and theta phase coherence.

## Results

LFPs were simultaneously recorded in the hippocampal pyramidal layers of CA3 and subiculum, hippocampal CA1 stratum radiatum, as well as ECoG in the parietal cortex (PCx) across vigilance states of eleven freely moving mice (see methods; Fig. 1A). We assessed the phase-amplitude coupling across a wide range of slow and fast oscillations using comodulogram analysis, which estimates coupling across different pairs of frequency bands (see methods). Individual comodulogram graphs were calculated for aWk, NREMs, and REMs from continuous 24 h LFP/ECoG recordings (Figs. 1A-1B, and S1). To investigate dynamics of coupling across multiple frequency and temporal scales, we computed the Modulation Index (MI)^17^ using a time-resolved approach (see methods).

### Theta modulates three distinct gamma bands during REMs

Using comodulogram analysis, we found that theta phase (6-9 Hz) significantly modulates middle gamma amplitude (MG: 50-90 Hz) across CA3, CA1, subiculum, and PCx recordings during aWk and REMs (p < 0.0001 for all, surrogate test). During REMs, theta-MG coupling strength was significantly increased for subiculum and PCx, but not for CA3 and CA1, recordings (CA3, CA1: p > 0.05; Sub: p < 0.01; PCx: p < 0.001; Wilcoxon test; n = 11 animals; Fig. 1D, Table S1). During REMs, in addition to theta-MG coupling, theta phase significantly modulated the ultrahigh gamma (UG: 160-250 Hz; p < 0.0001, surrogate test; Fig. 1B-1E) in pyramidal layers of CA3 and the subiculum, as well as high gamma (HG: 110-160 Hz; p < 0.0001, surrogate test) in the PCx. For all studied regions, theta-MG coupling was always stronger than theta-HG and theta-UG coupling during REMs (Fig. 1B). The comodulogram analysis of NREMs LFP recordings showed phase-amplitude coupling between different frequency pairs for different regions (Fig. S1). Ripples generated within the CA3 pyramidal region and CA1 stratum radiatum exhibited coupling to sharp waves (Fig. S1B). NREMs ripples from the pyramidal layer of subiculum were phase-coupled to both slow waves (0.5-1.5 Hz) and spindles (10-15 Hz).

We used phase-amplitude histograms obtained during calculation of the MI to study the preferred theta phases among MG, HG, and UG during aWk and REMs. We analysed CA3, subiculum and PCx recordings that showed distinct coupling patterns both in MG and either HG or UG across REMs (Fig. 1C). We found that the UG modulation significantly phase lagged behind MG modulation by ∼90° (∼32 ms time delay) in CA3 pyramidal layer, while no difference was observed for the subiculum pyramidal cell layer during REMs. For the PCx, a slight difference (∼20° or ∼8 ms time delay) was found between preferential theta phases of HG and MG. Furthermore, comparing preferred phases for MG during aWk and REMs revealed no difference (Fig. 1C).

**Figure 1.**
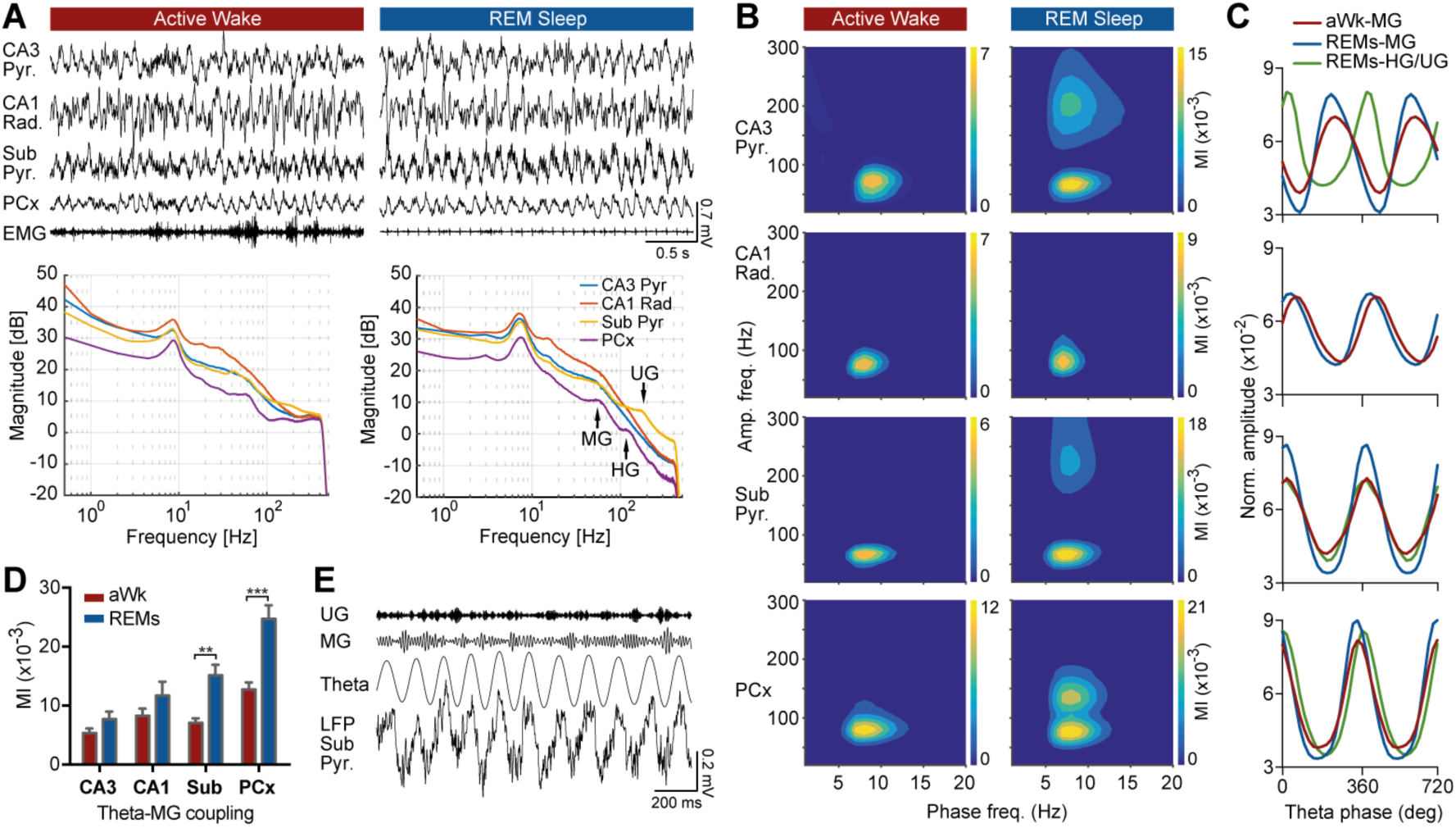
General patterns of phase-amplitude coupling during active wake and REMs. (*A*) Concurrent recordings from CA3 and subiculum pyramidal layers, CA1 radiatum, and PCx as well as EMG signal during aWk and REMs; corresponding power spectral density measurements obtained from 24 h recordings are also shown. Note two distinguishable peaks within the gamma range in the PCx and a peak in UG frequency range of subicular recordings during REMs. (*B*) Comodulogram graphs showing MI for a wide range of frequency band pairs, obtained from 24 h recordings of CA3 pyramidal layer, CA1 stratum radiatum, subiculum pyramidal layer, and PCx of one mouse. (*C*) Corresponding phase-amplitude histograms for the theta-MG and either theta-HG or theta-UG coupling during aWk and REMs for the same animal as in (*B*). The distributions are consistent across animals. Zero and 180 degrees correspond to trough and peak of theta cycle, respectively. (*D*) Quantification of theta-MG coupling during aWk and REMs (mean ±SEM; n = 11 animals; ** p < 0.01; *** p < 0.001; Wilcoxon test). REMs coupling is significantly stronger than aWk within the subiculum and PCx, but not within CA3 and CA1. (*E*) Example of UG in the subiculum pyramidal layer during REMs.

### Dynamics of theta-gamma coupling across REMs

Using a time-resolved approach, we investigated dynamics of the phase-amplitude coupling during sleep (see methods, Fig. 2A). We found that phase-amplitude coupling waxes and wanes over time during each REMs episode. Indeed, it is a discrete and transient phenomenon, which lasts for only few seconds with no detected periodicity. We also used the time-resolved phase-locking value (PLV) as a control measure of phase-amplitude coupling (Fig. 2B). Interestingly, this dynamical regulation of CFC is independent of the existence of gamma activity over the whole course of a REMs episode (Fig. 2C). Figure 2D shows dominant frequency of fast oscillation envelope, which was used as phase frequency to obtain time-resolved MI and PLV measures in Figs. 2A and 2B. To study the relation between coupling strength and theta/gamma power, we depicted these measures in a color-coded scatter plot (Fig. S3). Theta-MG coupling strength during aWk, specifically within CA3 and CA1, was less dependent on theta/gamma power than REMs (Fig. S3). During REMs, especially within subiculum and PCx, stronger theta-MG coupling is associated with higher theta/gamma power.

**Figure 2.**
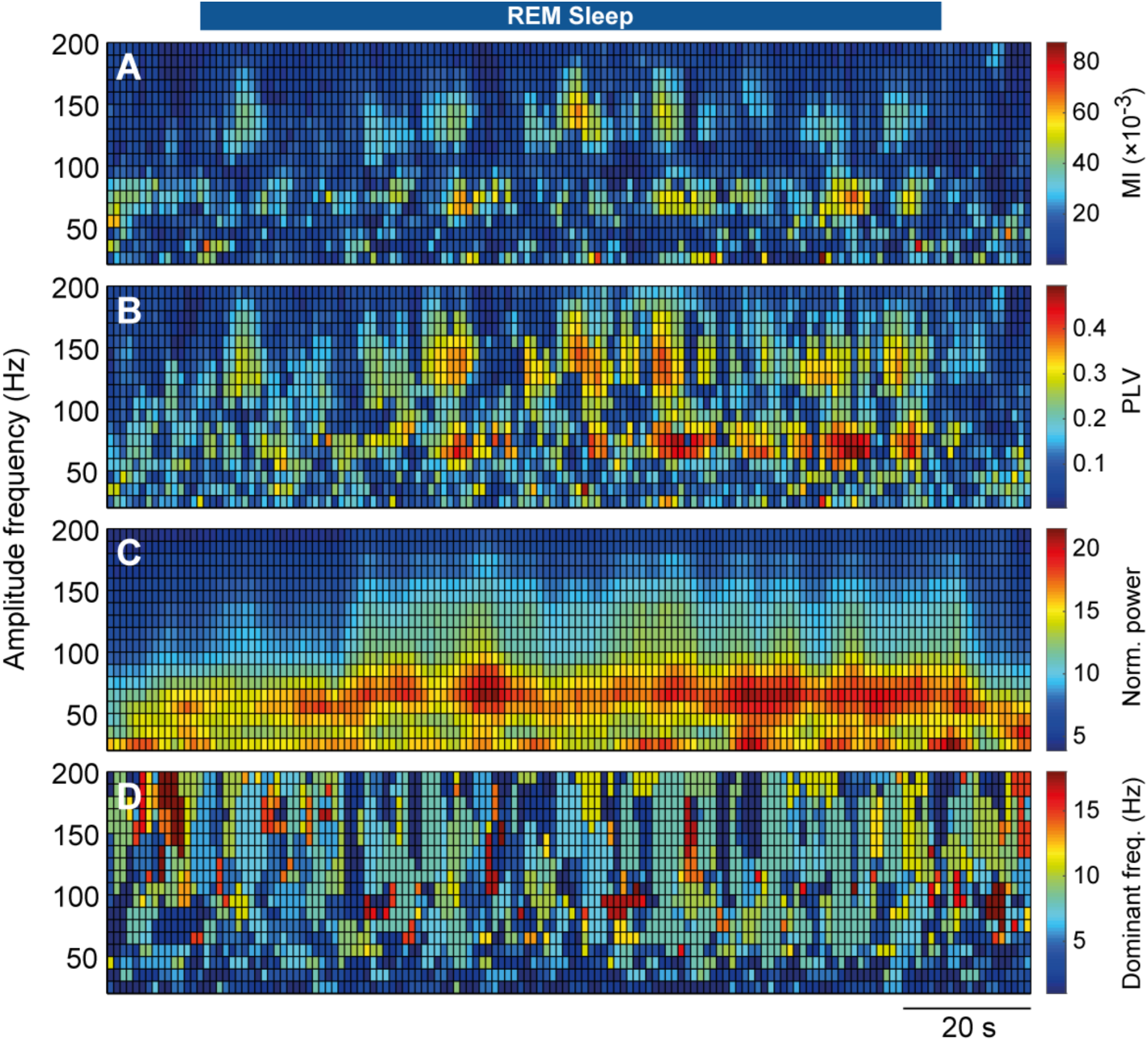
Dynamics of phase-amplitude coupling during REMs. (*A, B*) Time-resolved MI and PLV measures estimated from PCx across a REMs episode. Horizontal and vertical axes are time and modulated frequencies (i.e., gamma range). The measures were estimated using a 4-s moving window with 75% overlap. Blue bar in the top indicates the REMs episode. (*C*) Normalized power of fast oscillations obtained from 4 s moving windows (75% overlap) during a REMs episode. Power of amplitude frequency bands were normalized to their first 10 s average power, to provide meaningful comparison between frequency bands. (*D*) Dominant frequency of the envelope of fast oscillations, which is considered as phase frequency (modulating frequency) for the time-resolved approach.

### Theta-gamma coupling swings during REMs

The discrete nature of phase-amplitude coupling ignites the question as to what is happening in other hippocampal regions during off periods of coupling. To answer this, we studied theta-MG coupling changes simultaneously among CA3, CA1, subiculum, and PCx recordings within aWk and REMs episodes. Interestingly, we found that coupling swings between recorded hippocampal regions, while during some periods of time coupling exists concurrently between regions (Fig. 3A). We further investigated phase-amplitude coupling across CA1 layers using silicon probe recordings to further verify the existence of possible switching mechanisms between CA1 strata. We observed that theta-MG coupling is simultaneously occurring in all layers, without any switching between layers (Fig. 3B). To quantify concurrent coupling between regions, we estimated overlap ratio and correlation between pairs of channels during aWk and REMs (Figs. 3C-3D). Overlap ratio was restricted to periods of time where at least one region shows high levels of theta-MG coupling (1.5 SD + Mean), and indicates portion of these times where two regions show theta-MG coupling higher than 1.5 SD above the mean, simultaneously. We showed that overlap ratio is significantly increased during REMs in comparison to aWk for all studied pairs (p < 0.01, Wilcoxon test; Fig. 3C, Table S2), except for CA3-PCx. Correlation measure considers all periods of time during aWk or REMs, and is increased significantly during REMs in comparison to aWk for all studied pairs (p < 0.01, Wilcoxon test; Fig. 3D, Table S2). To examine the existence of periodicity in coupling events, we measured the distance between theta-MG coupling events (inter-coupling interval) within individual aWk and REMs episodes. We found that inter-coupling interval varies from a few seconds up to tens of seconds with an average ∼16 s, however, the average and distribution of inter-coupling interval is quite consistent across aWk and REMs (Figs. 3E and S4, Table S1).

**Figure 3.**
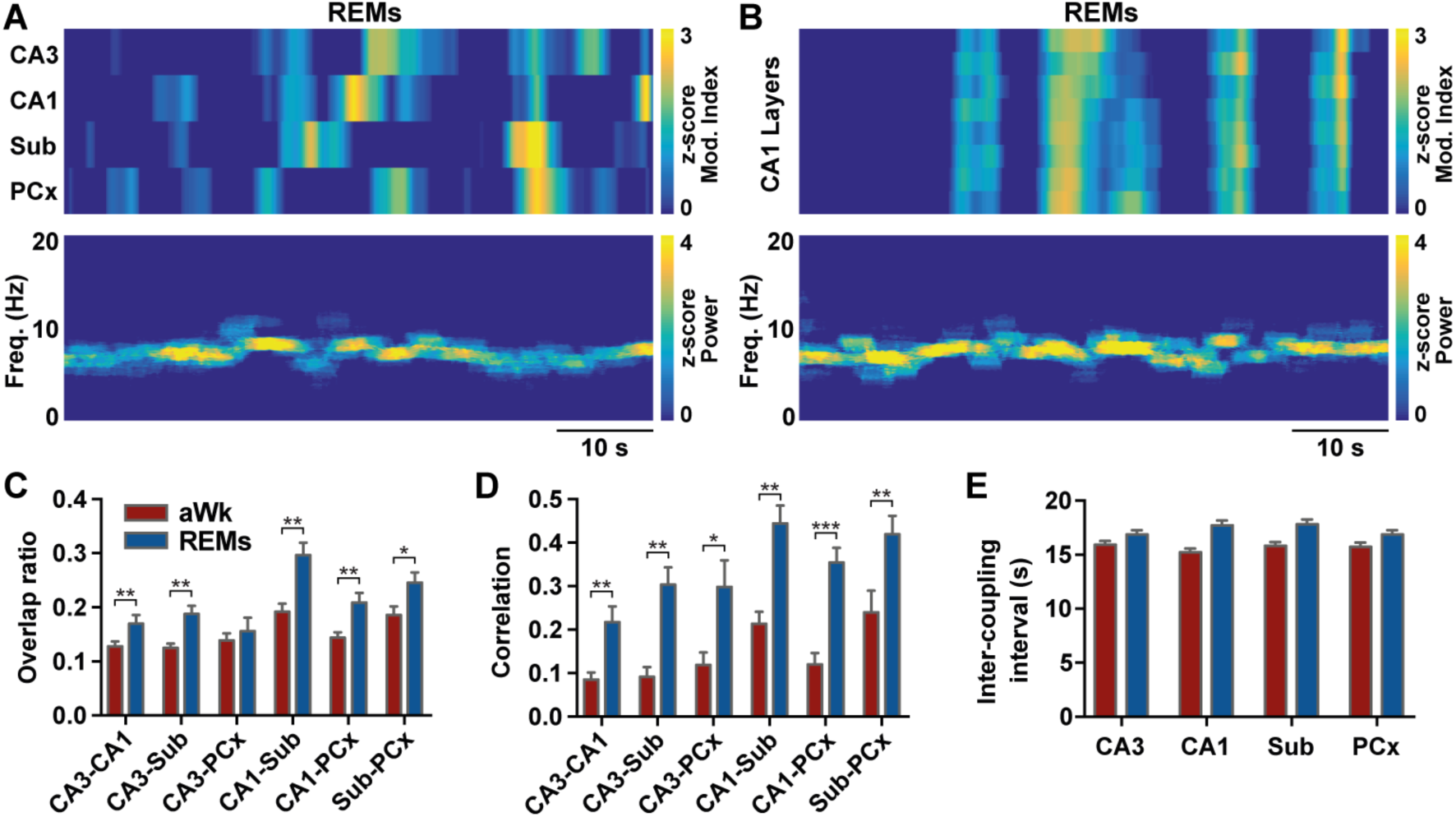
Theta-gamma coupling swings between hippocampal/cortical regions during REMs. (*A*) Top: dynamics of theta-MG coupling over a REMs episode for CA3, CA1, subiculum, and PCx recordings. Coupling switches between regions, but co-occurs during some time periods. The MI measure was estimated between theta and MG using a 4 s moving window with 75% overlap, and then smoothed by a 10-point Hanning window. Bottom: corresponding time-frequency representation. (*B*) Same as (*A*), except for different layers of CA1. Theta-MG coupling is highly synchronized across the layers of CA1. (*C*) Quantification of concurrency between recording pairs during aWk and REMs. Overlap ratio indicates the proportion of times that two regions simultaneously show theta-MG coupling higher than 1.5 SD above the mean. (*D*) Correlation between theta-MG coupling for different recording pairs. (*E*) Average distance between theta-MG coupling events. Inter-coupling interval indicates silent periods between two coupling events. All bar graphs represent the mean ±SEM values of 11 animals (* p < 0.05; ** p < 0.01; *** p < 0.001; Wilcoxon test).

### Theta-MG coupling increases during a single REMs episode

Temporal representations of coupling during individual REMs episodes consistently revealed that the intensity of coupling usually increases as an episode progresses (Figs. 2A and 3A). To further verify this finding, we divided each REMs event into 3 equal segments, and computed theta-MG coupling and theta/MG power for each segment. For this analysis, only REMs episodes which lasted more than 30 s were considered to provide segments with at least 10 s length. For each REMs episode, middle and late values were normalized to the early segment. We found that the theta-MG coupling significantly increased across REMs, for all studied regions (P<0.001 for all; Wilcoxon test; Fig. 4A, Table S3). No significant change in theta power across REMs was observed (P>0.05; Fig. 4B), while MG power significantly increased (p < 0.001 for all; Fig. 4C). We did not find any linear correlation between duration of REMs episode and MI changes, however shorter periods of REMs showed less change in coupling strength across the episode (Fig. 4D). In CA3 and CA1, average relative MI showed the highest increase for REMs episodes with ∼70 s duration, with decreasing values for longer REMs episodes in CA3, whereas remained stable in CA1 after ∼70 s. The MI changes in the subiculum and PCx were less dependent on duration of REMs.

**Figure 4.**
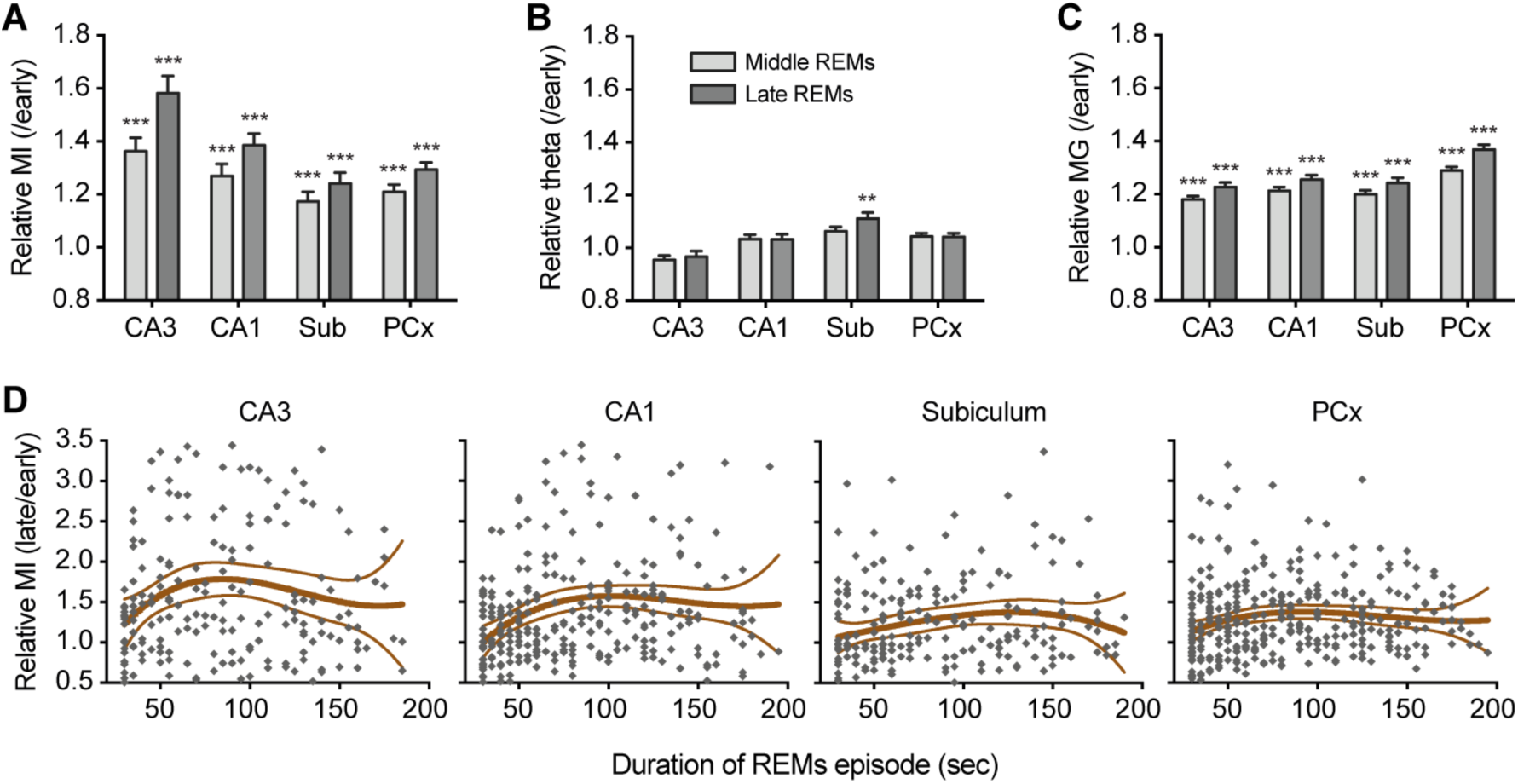
Theta-MG coupling strength increases over REMs. (*A*) Normalized MI values for CA3, CA1, subiculum, and PCx recordings during middle and late phases of REMs episodes (mean ±SEM; n = 11 animals; *** p < 0.001; Wilcoxon test). For each REMs episode, middle and late MI values were normalized to the early segment. Theta-MG coupling increased significantly for all studied regions. (*B, C*) Same graphs as (*A*) for theta and MG power, respectively. There were no significant changes in theta power across REMs segments (except for the subiculum; ** p < 0.01), while MG power increased significantly. (*D*) Relation between duration of REMs episodes and the change in theta-MG coupling for studied regions. Each point represents one REMs episode. Lines indicate mean ±SEM.

### Optogenetic silencing of MS^GABA^ neurons suppresses theta-gamma coupling and theta phase coherence

In a previous study^3^, we showed that optogenetic inhibition of MS^GABA^ cells during REMs impairs object recognition and fear-conditioned contextual memory in mice. To investigate a possible underlying mechanism, we calculated power spectral density, theta phase coherence, and phase-amplitude coupling of hippocampal/cortical recordings before, during, and after optogenetic silencing of MS^GABA^ neurons within REMs (Fig. 5, Table S4). Consistent with our previous study^3^, we found that although theta power is significantly reduced during inhibition (p < 0.001 for all recording sites; n = 5 animals; Figs. 5A and 5F), the MG/HG/UG power show no significant changes (p > 0.05 for all recording sites and gamma bands; Fig. 5F). Importantly, here we found that optogenetic silencing of MS^GABA^ neurons significantly decreased both theta phase coherence and theta-gamma coupling within the hippocampus and PCx (Figs. 5B-5F). There was a strong phase coherence in the theta range between recorded sites before and after stimulation, while it was significantly decreased during MS^GABA^ optogenetic silencing (p < 0.001 for all recording pairs; Figs. 5B-5D). The theta-gamma coupling strength was also significantly decreased during silencing of MS^GABA^ cells (p < 0.001 for all recording sites and gamma bands; Figs. 5E-5F).

**Figure 5.**
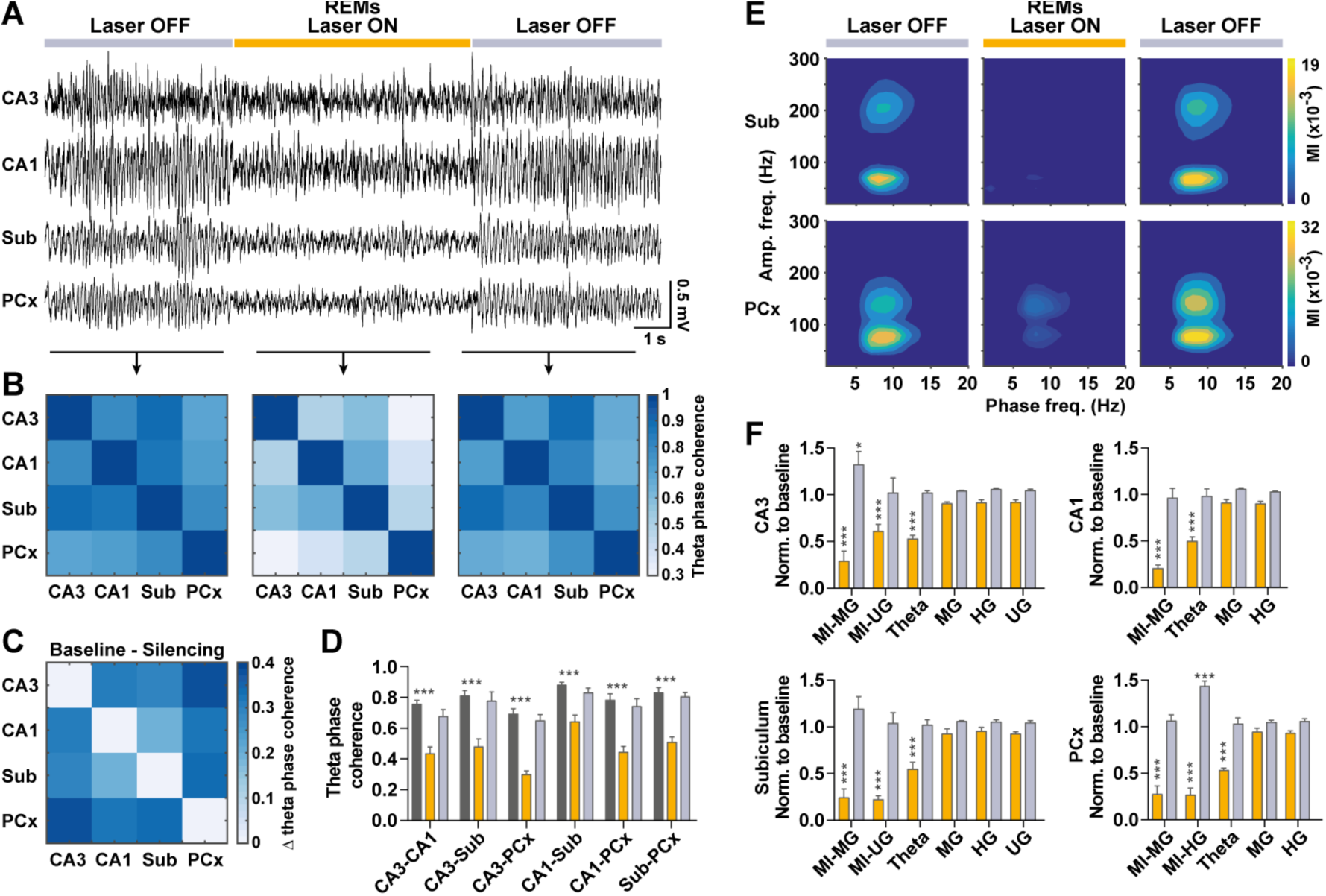
Optogenetic suppression of extrinsic theta projections disrupts coupling and theta phase coherence. (*A*) REMs hippocampal and cortical recordings before, during, and after optogenetic inhibition of MS^GABA^ neurons. Baseline was considered as 5 s of REMs immediately before silencing. (*B*) Theta phase coherence between CA3, CA1, subiculum, and PCx recording sites, before/during/after inhibition. Graphs show average results of 50 inhibition trials from 5 mice. There was a strong phase coherence between recorded sites before and after inhibition, while coherence was significantly decreased during MS^GABA^ inhibition. (*C*) Difference in theta phase coherence between baseline and silencing. The highest decrease in coherency was between CA3 and PCx sites, which dropped from ∼0.7 to ∼0.3. (*D*) Quantification of theta phase coherence before/during/after inhibition, which is significantly decreased during silencing in comparison to the baseline for all recording pairs. (*E*) Coupling comodulograms before/during/after MS^GABA^ inhibition estimated from 10 trials of one mouse. (*F*) Changes in theta-MG (MI-MG), theta-HG (MI-HG), and theta-UG (MI-UG) coupling, theta power, and middle/high/ultrahigh gamma power, during and after inhibition in comparison to baseline. All reported values in (*D, F*) are mean ±SEM, obtained from 5 animals, 10 trials each. There is no difference between vertical and horizontal asterisks (* p < 0.05; *** p < 0.001; Wilcoxon test).

## Discussion

Existence of coupling in a network demonstrates a functionally active circuit^34^. On the other hand, for proper information processing among different brain regions, each local network needs to integrate its input/output at a given time. That is, information processing is distributed both in time and space, where theta waves may represent a possible underlying orchestrator of information processing during sleep through phase coding. Therefore, the orchestrated nature of theta-gamma coupling may provide necessary mechanisms for communication between different brain regions while limiting interference, and supports our hypotheses of specific phase-space coding of information processes during REMs. More specifically, the off periods in the coupling graph of one recording region may indicate those times when the other regional networks are turned on. In our analysis, we have considered four brain networks of hippocampal CA3, CA1, subiculum and PCx, and noticed that coupling predominantly swings between them, while co-occurring for relatively small time periods. However, we found time periods where significant coupling was simultaneously absent from all recorded regions, which suggests that such circumstances might correspond to the time periods when other brain areas, such as other cortical or thalamic sites, are integrating their input/output.

The time periods that pairwise combination of recording sites exhibited high levels of coupling, simultaneously, were significantly increased during REMs in comparison to aWk. We measured coupling concurrency of two recorded sites using overlap ratio and correlation; overlap ratio considers only segments with high level of coupling (1.5 SD + Mean), while correlation is obtained from whole segments. Both measures significantly increased for all pairwise combinations (except overlap ratio between CA3 and PCx), and highest concurrency during REMs was observed between CA1 and subiculum and then between subiculum and PCx. This indicates that two networks are activated together during REMs more frequently than aWk, which provides further windows of opportunity for communication and information transfer between networks during REMs. In addition, the inter-coupling interval histograms showed same distributions for aWk and REMs (Figs. S4), indicating that increase in concurrent coupling during REMs is due to the way in which theta-MG coupling is orchestrated among recording sites, and not simply a result of an increased number of coupling events. Altered, and likely increased, patterns of coupling concurrency may be involved in the development of epileptic seizures in rodents considering that more than 80% of seizures are initiated during aWk (34%) and REMs (47%) in rats^35^, an observation which warrants further investigation. Finally, the results of silicon probe recordings demonstrated that theta-MG coupling occurs simultaneously across all CA1 layers. This suggests that hippocampal CA1 strata build an internal network and are uniformly activated during the presence of theta-MG coupling and deactivated during the absence of coupling.

### Subicular and cortical theta-MG coupling is significantly increased during REMs

The subiculum is the primary output of the hippocampus, as subicular pyramidal cells send projections to many brain structures including cortical, thalamic, hypothalamic, and septal nuclei^36,37^. Regular-spiking and bursting cells are two distinct classes of subicular pyramidal neurons with possibly different functions^38,39^. Subicular UG, which is important for robust information transfer^40^, is generated by the activation of bursting cells while regular-spiking neurons are inhibited^39,41^. Considering phase-amplitude coupling as a mechanism for spatial information processing and transmission^26,42^, insignificant level of theta-UG coupling during aWk (p > 0.05, surrogate test) may indicate a lack of spatial information transfer to neocortical sites for more permanent storage. However, during REMs, the existence of such coupling promotes routing information out of the hippocampus to neocortical sites for long-term memory storage.

Interestingly, *in vitro* work has also demonstrated that subicular theta-UG coupling was significantly decreased in a transgenic mouse model of Alzheimer’s disease^26^, which highlights the potential necessity of theta-UG coupling for forming new memories. Furthermore, increased theta-MG coupling in subicular and cortical, but not CA3 and CA1, sites during REMs in comparison to aWk, indicates that the subicular and cortical structures serve critical roles as hippocampal output and long-term memory storage target, respectively, in hippocampal-neocortical dialog.

Existence of independent theta nested gamma oscillations has been hypothesized to provide the basic mechanism for memory retrieval and encoding^43,44^, while the occurrence of distinct gamma bands on different theta phases may help to minimize the interference between retrieval and encoding streams^44,45^. Our findings thus support the hypothesis that memory encoding and retrieval is mediated through modulation of specific gamma bands, however their preferred modulation phase may not play a role, as our findings demonstrated that among three recording regions that showed two distinct phase-locked fast oscillations, only in recordings from the CA3 pyramidal layer are MG and UG oscillations modulated at significantly different phases of theta cycle (90° ≈ 32 ms). No preferred theta phase difference was observed for MG and UG for recordings completed at the subicular pyramidal layer, whereas for the PCx only a slight difference (∼20° or ∼8 ms time delay) was found between preferential theta phases of HG and MG, which is negligible relative to the duration of a theta cycle (∼140 ms).

In contrast to aWk and REMs, slow waves, spindles, and hippocampal sharp wave-ripples (SWRs) are prevalent during NREMs, which are temporally coupled and facilitate hippocampal-neocortical dialog^46-49^. Hippocampal SWRs are associated with the replay of synaptic activities critical for the encoding of space/context/time^50^, and facilitating information transfer within hippocampal-cortical circuits^51,52^. SWRs involve cycles of 50-150 ms bursting spikes, which propagate along the CA3-CA1-subiculum axis^53^. Coupling between ripples and spindles in human has been previously reported in hippocampal^54^ and parahippocampal sites^46^. In line with previous observation^54^, our findings show that hippocampal ripples are also phase-locked to both spindles and slow oscillations within the subiculum in mice, although for CA3 and CA1 recording sites ripples were only coupled to sharp waves (Fig. S1). This suggests that the coordination of spindles with ripple oscillations in the subiculum could be a primary mechanism mediating hippocampal-neocortical dialog during NREMs.

### Both middle gamma and theta-MG coupling strengthened across REMs

In humans, gamma oscillation during REMs is correlated with dream lucidity, where gamma power during lucid REMs is significantly higher than during non-lucid REMs^55,56^. Furthermore, transcranial alternating current stimulation at gamma frequency, but not other frequency bands, induced conscious awareness during REMs and increased dream lucidity in humans^57^. A recent study also showed correlation between dream content and region-specific gamma activities during REMs in humans^58^. These studies suggest that the increasing gamma and theta-MG coupling across REMs may progressively reinforce conscious awareness, eventually lead to an arousal. This mechanism can be further supported by comparing gamma power during aWk and REMs, where average MG power during REMs was lower than aWk (Fig. 1A), and MG power within individual 4 s segments of aWk and REMs overlap to some extent (Fig. S3). Therefore, a fuzzy thresholding in either MG power or theta-MG coupling level may be responsible for behavioural state switching. This remains speculative at this point and requires further investigation.Our results also showed that although there is no linear correlation between duration of REMs episodes and changes in theta-MG coupling, these changes do tend to be less in shorter REMs episodes. On average, the highest increase in theta-MG coupling occurred in REMs episodes with ∼70 s in CA3, CA1, and PCx recording sites. Interestingly, the average REMs duration in mice is around 60-70 s, which might indicate an optimal duration for gamma power and theta-MG coupling to reach their highest levels before transition to wakefulness.

### Theta REMs is essential to maintain theta-gamma coupling and theta phase coherence

*In vivo*, the majority of hippocampal theta waves are generated by extrinsic projections originating from the medial septum^3,31,59^. Optogenetic silencing of MS^GABA^ neurons significantly decreased theta power, theta phase coherence, and phase-amplitude coupling by ∼55%, ∼40%, and ∼75%, respectively. However, the power of fast oscillations remained nearly unaltered (∼10% reduction). Gamma oscillations is generated by monosynaptic closed-loop interaction between excitatory pyramidal cells and fast-spiking inhibitory interneurons, and their power is independent of hippocampal theta power^3,34^. Therefore, the hippocampal theta rhythm may facilitate the organization of gamma oscillations without the large-scale activation of place cells. However, the power of theta oscillations influences both theta phase coherence and theta-gamma coupling strength, highlighting the crucial role of theta power, and not only theta phase, in providing the mechanisms necessary for information processing.

The “communication through coherence” hypothesis suggests that two brain networks should oscillate coherently during efficient information transfer^60,61^. Indeed, phase coherence facilitates neural communication and spike-timing dependent plasticity, which each contribute to long-term memory formation^62,63^. On the other hand, phase-amplitude coupling contributes mainly to short-term memory formation^64^ and facilitates non-interfering multiple object representation in working memory^62,65^. By optogenetically suppressing the theta rhythm, which has been shown previously to impair contextual memory ^3^, we indeed impeded these two mechanisms essential for working and long-term memory.

## Methods

### Animals

Male VGAT-ires-Cre (VGAT::Cre) transgenic mice (The Jackson Laboratory, stock number 016962) were used. All mice were housed individually in polycarbonate cages and kept at constant temperature (22° C) and humidity (30-50%) with a constant light cycle (12 h : 12 h, light : dark; lights on at 0800). Mice were allowed unlimited access to food and water. Animals were treated according to protocols and guidelines approved by McGill University and the Canadian Council of Animal Care.

### Virus-mediated targeting of opsin and eYFP expression

At ∼10 weeks age, mice were anesthetized with isoflurane (5% induction, 1-2% maintenance) and placed in a stereotaxic frame. Recombinant AAVdj (0.6 μL) was injected through a 28 G cannula into the medial septum (MS) at the following coordinates (in mm, relative to bregma); anterior-posterior (AP) +0.86; medial-lateral (ML) 0.0; dorsal-ventral (DV) -4.5. An in-depth evaluation of the precision and efficacy of this strategy for enabling ArchT-eYFP-mediated inhibition of MS^GABA^ neurons was completed in a prior study^3^.

### Electrode and optic fiber implantation for in vivo experiments

Tetrodes were made by twisting four individual 17.5 μm diameter platinum-iridium (platinum:iridium 90%:10%) wires into a single strand. For targeting dorsal hippocampal areas CA3, CA1 and subiculum, leads from 3 tetrodes were then soldered to an electrode interface board and fixed in the correct orientation using epoxy to facilitate implantation as a single array. Tetrode ends were cut to the appropriate length in relation to one another and subsequently cleaned immediately prior to surgery, giving a measured impedance of ∼1 MΩ.

At 18 weeks age mice were anesthetized with isoflurane (5% induction, 0.5-2% maintenance) and subsequently placed in a stereotaxic frame. After clearing the skull of connective tissue and drying with alcohol, holes were drilled in the bone above the dorsal hippocampus (CA3: AP -1.8, ML +2.1; CA1: AP -2.45, ML +1.8; subiculum: AP -3.10, ML +1.5; Fig. S5). The dura was gently cut, and the electrode array was slowly lowered until the tips of the tetrodes in the pre-arranged array were at the correct depth (relative to Bregma; CA3: DV -2.25; CA1: DV -1.3; subiculum: DV -1.5; Fig. S5). A similar procedure was used for implantation of silicon probes (NeuroNexus, Ann Arbor, MI, USA) used to record laminar local field potentials at 50 μm increments oriented vertically through hippocampal CA1, although slightly different coordinates were used for optimal orientation of the probe perpendicular to the CA1 layers (AP -2.45, ML +1.5, probe tip DV -1.5). In all experiments, a screw implanted in the skull above the parietal cortex (AP, -2.3; ML, -1.35) that is contacting the surface of brain tissue served as ECoG, while 2 stranded tungsten wires inserted into the neck musculature were used for EMG recording for assessing postural tone. Screws placed in the bone above the frontal cortex and cerebellum served as ground and reference, respectively. An optic fiber implant, used to deliver laser light to the medial septum virus-transfection zone for optogenetic experiments completed on the same mice for a different study, was also implanted just above the medial septum (AP, +0.86; ML, -0.2; DV, -3.82; Fig. S5). As a final step, dental cement was applied to the skull to permanently secure all components of the implant.

### In vivo electrophysiological recording and optogenetic MS^GABA^ inhibition

Mice were allowed to recover from surgery undisturbed for at least 1 week. Afterwards, a headstage preamplifier tether was attached to a connector on the top of the implanted electrode interface board. Mice were subsequently returned to their home cage where they were allowed to habituate to the tether system for 1 week. Once mice were habituated to the chronic tethering system, and before any optogenetics experiments commenced, a 24 h consecutive baseline recording was completed. Following baseline recording, an additional recording session ∼2 h in duration was conducted to assess how disrupting MS^GABA^ activity would influence cross-frequency coupling and theta phase coherence during REMs. Mice were continuously monitored until they entered into at least 10 s of stable REMs (see criteria in ‘Determination of vigilance state’ below), at which time continuous square pulses of orange light, ∼7 s in duration, were delivered to the MS transfection zone via the optic fiber implant at a frequency of ∼2 pulses/minute. This protocol was continued until mice transitioned to wakefulness, at which point MS^GABA^ inhibition ceased until subsequent REMs occurred. The intensity of light delivered to the MS was calibrated to ∼20 mW, a value that, in the absence of ArchT expression, was previously shown to have no effect on baseline EEG characteristics and animal behavior ^3^. The precise onset and offset of light delivery was timestamped to ECoG/LFP/EMG recordings. For each session, all recorded signals from implanted electrodes were amplified by the headstage preamplifier tether before being digitized at 16000 Hz using a digital recording system (Neuralynx, Inc.) and saved to a hard disk. After applying a low pass filter (Chebyshev Type I, order 8, low pass edge frequency of 400 Hz, passband ripple of 0.05 dB) to prevent aliasing, all recordings were downsampled to 1000 Hz to obtain LFP signals.

### Histological confirmation of electrode/optic fiber placement and construct expression

Following completion of all experiments, mice were deeply anesthetized with ketamine/xylazine/acepromazide (100, 16, and 3 mg/kg, respectively, intraperitoneal injection) and electrode sites were subsequently marked by passing a current of 10 μA for ∼10 s through each of the tetrodes. Following lesioning, mice were perfused transcardially with 1 x PBS-heparine 0.1%, pH 7.4, followed by 4% paraformaldehyde in PBS (PFA). Brains were removed from the skull and placed in PFA at 4 ° C overnight before being cryoprotected in 30% sucrose dissolved in PBS at 4 ° C for an additional 24 h. Brains were then sectioned (50 μm) using a cryostat; half of sections collected were further processed to evaluate accuracy of ArchT-eYFP construct expression (see paragraph below), whereas the remaining sections were mounted on gelatin-coated glass slides, stained with cresyl violet and coverslipped. Final electrode and optic fiber locations were determined by viewing these stained sections with a light microscope. Only mice with confirmed placement of electrodes in the CA3 pyramidal layer, CA1 stratum radiatum (all layers for silicon probe-implanted mice), and the subiculum pyramidal layer, as well as proper optic fiber tip placement immediately above the MS transfection zone, were further analysed.

To confirm accurate targeting of the ArchT-eYFP construct to MS^GABA^, sections not used for electrode / optic fiber localization were initially washed in PBS 1x-Triton (0.3%) (PBST). Sections were then incubated for 60 minutes at room temperature in a blocking solution composed of 4% bovine serum albumin (BSA) dissolved in PBST prior to being incubated in rabbit anti-GFP (diluted 1:5000 in BSA) overnight at 4° C. To detect the primary antibody, sections were incubated the following day in alexa fluor 488 ex anti-rabbit IgG (H+L) (diluted 1:1000 in PBST) for 1 h at room temperature, prior to being mounted onto glass slides and coverslipped with Fluoromount-G. Once slides were sufficiently dry, fluorescent images of immunolabelled sections were acquired using a fluorescent microscope. Only mice with confirmed construct expression restricted to neurons localized within the MS and diagonal band regions were further analysed.

### Data analysis

All simulations and analyses were performed in MATLAB^®^ (R2016a, Natick, MA, USA) environment using custom scripts and built-in functions.

### Determination of vigilance state

Vigilance state was manually scored in 5 s epochs for all recordings through the concurrent evaluation of ECoG/LFP characteristics, EMG-derived muscle activity and video monitoring of behaviour. Quiet wakefulness (qWk) was defined as awake periods that mice were immobile including feeding and grooming behaviours, associated with absence of theta band ECoG/LFP activity and tonic muscle activity. Active wakefulness was defined as periods of theta band ECoG/LFP activity and EMG bursts of movement-related activity. NREMs was identified as periods with a relatively high amplitude low-frequency ECoG/LFP and reduced muscle tone relative to wakefulness associated with behavioural quiescence. REMs was defined as sustained periods of theta band ECoG/LFP activity and behavioural quiescence associated with muscle atonia, save for brief phasic muscle twitches. Polysomnographic scorings were performed double-blinded and was statistically confirmed to lie within a 95% confidence interval^66^. Statistics of vigilance states including percentage, duration, and density are provided in Fig. S6.

### Modulation Index (MI)

Phase-amplitude coupling was measured using Modulation Index (MI)^17^. To obtain MI, LFP/ECoG signals were first bandpass filtered into desired low and high frequency bands. The instantaneous phase of low frequency waveforms and the envelope of high frequency variations was subsequently estimated using the Hilbert transform. Afterwards, phase of the low frequency was discretized into several bins of equal sizes (*N* = 18 bins, each bin as 20°), and average power of the fast oscillations’ envelope was calculated inside each phase bin. The resulting phase-amplitude histogram (*P*) was compared with a uniform distribution (*U*) using Kullback-Leibler distance (*D*_*KL*_),

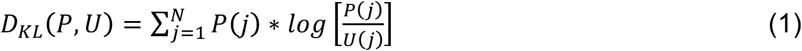

which in turn was normalized by *log(N)* to obtain MI (*MI = D*_*KL*_ */ log(N)*). LFP/ECoG signals were bandpass filtered using finite impulse response (FIR) filters in the both forward and revers directions to eliminated phase distortion (filtfilt, MATLAB signal processing toolbox). FIR filters were designed using the window-based approach (fir1, MATLAB signal processing toolbox), with an order equal to three cycles of the low cutoff frequency.

### Comodulogram analysis

Comodulogram analysis, which is a data-driven approach to explore coupling across different pairs of frequency bands, was used to assess the coupling between the phase of low frequencies and the power of high frequencies. We considered 18 frequency bands for phases from 0.5–18.5 Hz through 1 Hz increments and 2 Hz bandwidth, and 28 frequency bands for amplitude from the 20–310 Hz range through 10 Hz increments having 20 Hz bandwidth. MI values were then calculated for any of these pairs to obtain a comodulogram graph. For each mouse, we first filtered continuous 24 h recordings into the mentioned frequency bands and estimated the Hilbert transform, and then concatenated episodes of each vigilance state to derive the stage specific comodulogram graphs. To avoid power line interferences, those frequency bands in the vicinity of 60 Hz and its harmonics were re-assigned with a 2 Hz safe margin from these interfering frequencies. In comodulogram graphs, warm colors indicate existence of coupling between corresponding frequencies as indicated by the horizontal and vertical axes. Since MI is independent of absolute power of amplitude frequency, the resulting comodulogram graph is not biased for frequency bands with larger power. Therefore, MI based comodulogram analysis provides a powerful tool to assess phase-amplitude coupling for a wide range of frequencies.

### Time-resolved phase-amplitude coupling

A time-resolved coupling approach was investigated to track changes in phase-amplitude coupling over time and frequency^67^. The logic behind this approach is that if there exists a phase-amplitude coupling between low and high frequency bands, the modulating low frequency and the dominant frequency of the modulated gamma envelope should be identical. Therefore, by replacing frequency for phase axis in the comodulogram (horizontal axis) with the time axis, variations of phase-amplitude coupling over time and frequency can be followed. In short, the LFP/ECoG signal was first bandpass filtered into 24 sub-bands for amplitude frequency, as described previously for comodulogram analysis, and the envelopes of filtered signals are obtained using the Hilbert transform. The resulting envelopes were segmented into 4 sec windows with 75% overlap, and fast Fourier transform (FFT) of envelopes was calculated for each segment. To minimize the side effects of filtering on signal edges, we applied segmentation after filtering. The specific frequency for phase was obtained from the dominant frequency in the envelope of the filtered signal amplitude. The peak frequency of the envelope was found using the FFT, and the LFP/ECoG signal was bandpass-filtered at this peak frequency using a 2 Hz bandwidth. MI was then calculated as described above between the phase of this frequency band and the corresponding high frequency band.

### Theta phase coherence

Theta phase coherence between two signals was estimated using mean phase coherence^68^, which is the most prominent measure of phase synchronization. To obtain theta phase coherence, raw LFP/ECoG traces were first filtered in the theta range (6-10 Hz) using a 500^th^-order FIR filter (fir1, MATLAB) in both the forward and reverse directions. Afterward, the instantaneous phases of filtered signals were estimated using the Hilbert transform, and theta phase coherence was obtained by averaging the instantaneous phase differences of two filtered signals projected onto a unit circle in the complex plane. This measure has a value between [0 1], where zero and one indicate completely incoherent and coherent theta rhythms, respectively.

### Spectral analysis

Power spectral density (PSD) was estimated using the Welch’s method (pwelch, MATLAB signal processing toolbox), using 4-s segments with 75% overlap. Simply, the Welch’s method applies a Hamming window on each segment, calculates the FFT of the segments, and then averages over all resulting FFTs. Time-frequency representations were obtained using multitaper method from Chronux signal processing toolbox (window size = 3 s, step size = 0.1 s, tapers [3 5]).

### Statistical methods

The reported values are mean with the standard error of the mean (mean ±SEM). Two-tailed Wilcoxon signed-rank test was used for all reported statistics, unless otherwise mentioned. To evaluate phase-amplitude coupling results, surrogate data testing was performed to find out whether the MI values were statistically significant. Surrogate data were generated by shuffling the phase of the low frequency band 200 times, and MI values were calculated between these shuffled phase time series and the envelope of the amplitude frequency. Reported *p*-values were calculated by dividing the number of surrogate MI values greater than the original MI value by 200.

## Acknowledgments

We thank the Tidis laboratory members for insightful comments and discussions on previous versions of this manuscript. M.B. and K.S. were supported by Inselspital University Hospital. A.A. was supported by the Human Frontier Science Program (RGY0076/2012), Inselspital University Hospital, Swiss National Science Foundation and the University of Bern.

## Author contributions

M.B., K.S., and A.A. designed the analysis of data collected by R.B.;

M.B. analysed data; All authors discussed the results. M.B., R.B., and A.A. wrote the manuscript with contributions from all authors.

## Supplementary Information

**Figure S1.**
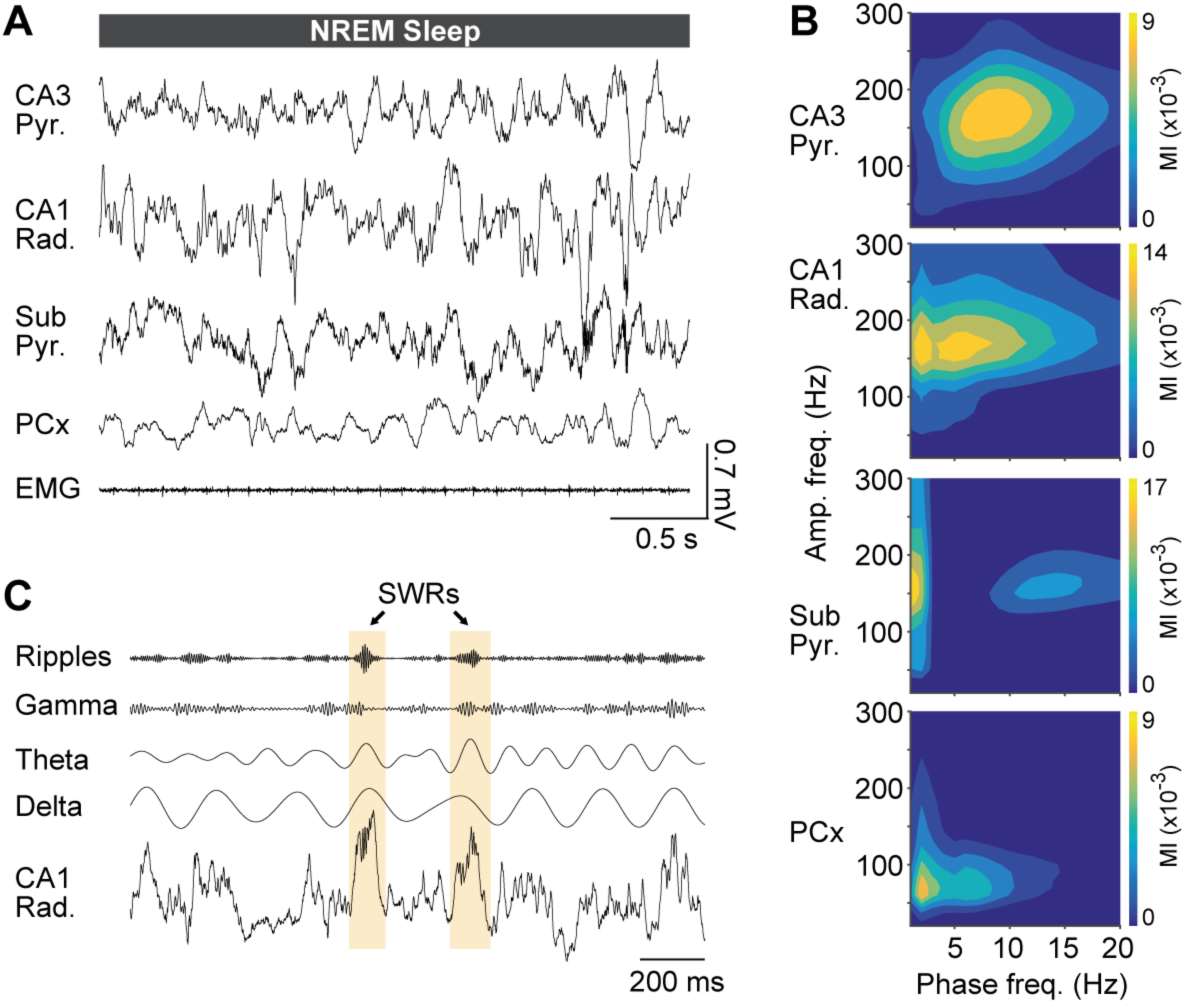
General patterns of phase-amplitude coupling during NREMs. (*A*) Concurrent recordings from CA3 pyramidal, CA1 radiatum, subiculum pyramidal strata, PCx and EMG signals during NREMs. (*B*) Comodulogram graphs show MI for a wide range of frequency band pairs, obtained from NREMs episodes of 24 h recordings performed from CA3 pyramidal layer, CA1 stratum radiatum, subiculum pyramidal layer, and PCx sites. (*C*) Example of sharp waves and their associated ripples (SWRs) in CA1 radiatum. As sharp wave spectra spread over a wide range of frequencies depending on SWRs duration, comodulogram graphs showed significant widespread slow modulating frequencies for the range of 2-15 Hz.

**Figure S2.**
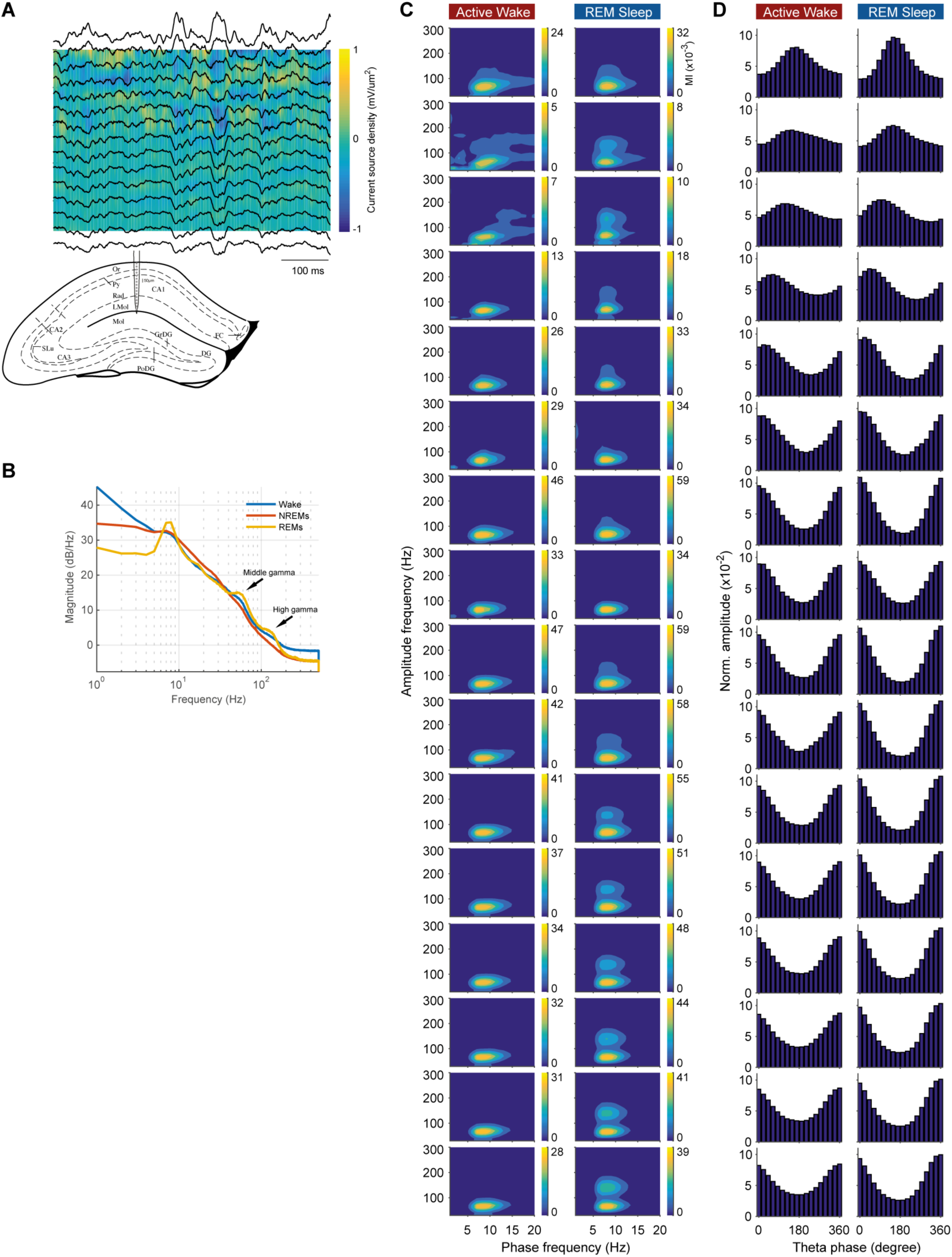
General patterns of phase-amplitude coupling across CA1 strata. (*A*) Current source density analysis and schematic of implanted probe. CA1 strata were recorded at 50 μm increments using a 16-channel linear probe. (*B*) Power spectral density during wake, NREMs, and REMs. (*C*) Comodulograms show phase-amplitude coupling across CA1 strata during vigilance states estimated from raw traces. (*D*) Phase-amplitude histograms of frequency pairs with highest coupling level in (*C*).

**Figure S3.**
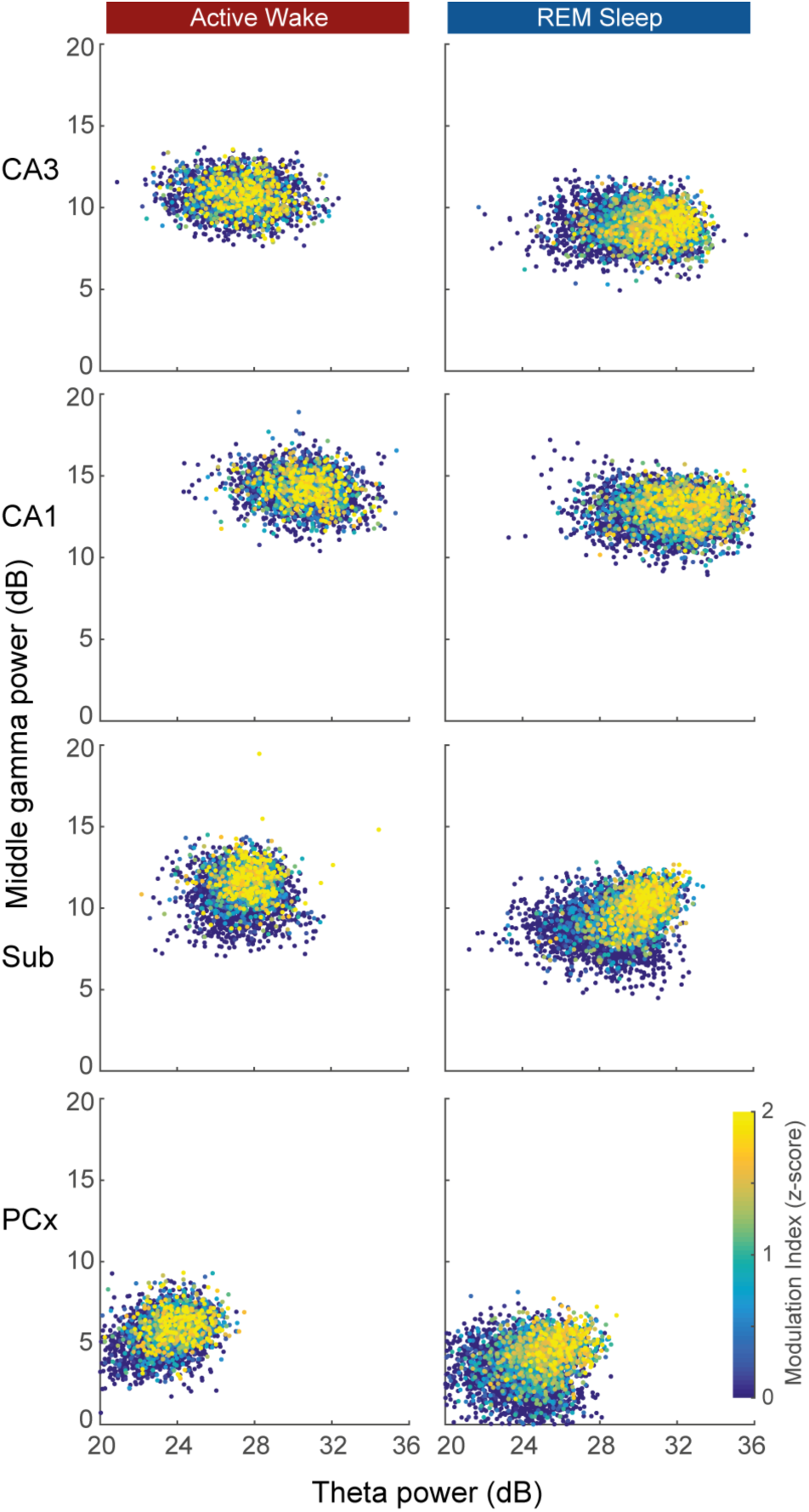
Relation between theta power, middle gamma power, and coupling strength. Scatter plots show distribution of theta and MG power during aWk and REMs and their corresponding theta-MG coupling strength, for CA3, CA1, subiculum, and PCx recordings. Each point represents a 4-s window, with consecutive windows having 75% overlap. Color code is normalized theta-MG coupling, where warmer colors indicate stronger levels of coupling. Theta-MG coupling strength during aWk, specifically within CA3 and CA1, was less dependent on theta/MG power than REMs. During REMs, especially within subiculum and PCx, stronger theta-MG coupling is associated with higher theta/MG power.

**Figure S4.**
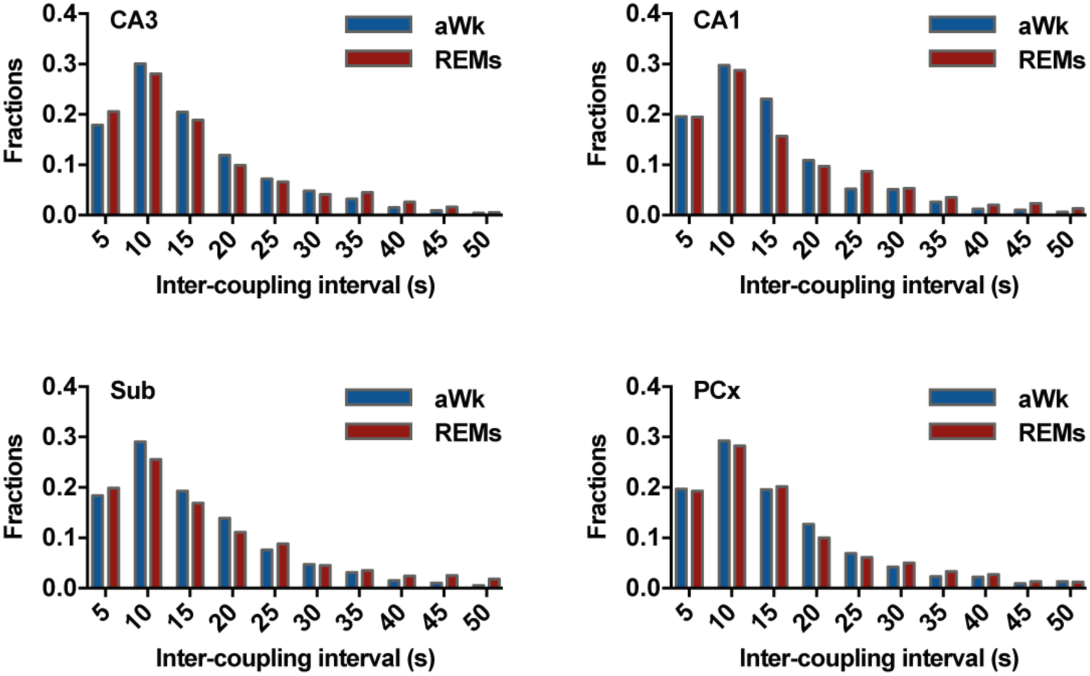
Inter-coupling interval histograms during aWk and REMs. Histograms show inter-coupling interval for CA3, CA1, subiculum, and PCx recordings for both aWk and REMs. There was no significant difference between aWk and REMs distributions (median values: CA3: aWk: 13 s; REMs: 12.8 s; CA1: aWk: 12.6 s; REMs: 13 s; Sub: aWk: 13.2 s; REMs: 13.4 s; PCx: aWk: 12.8 s; REMs: 13 s).

**Figure S5.**
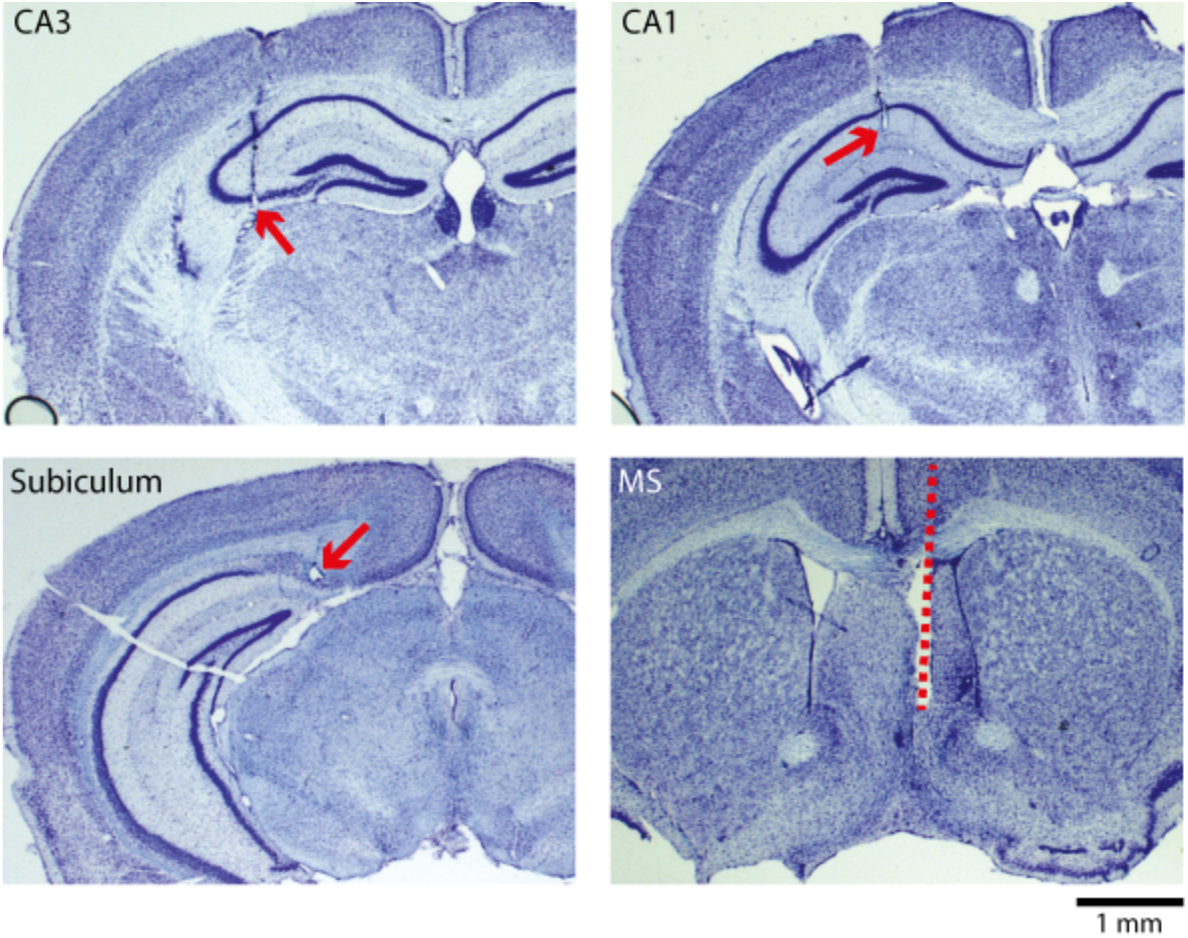
Placement of tetrodes and MS optic fiber. Sample coronal sections from tetrodes implanted in CA3, CA1, and subiculum, as well as optic fiber implanted in medial septum.

**Figure S6.**
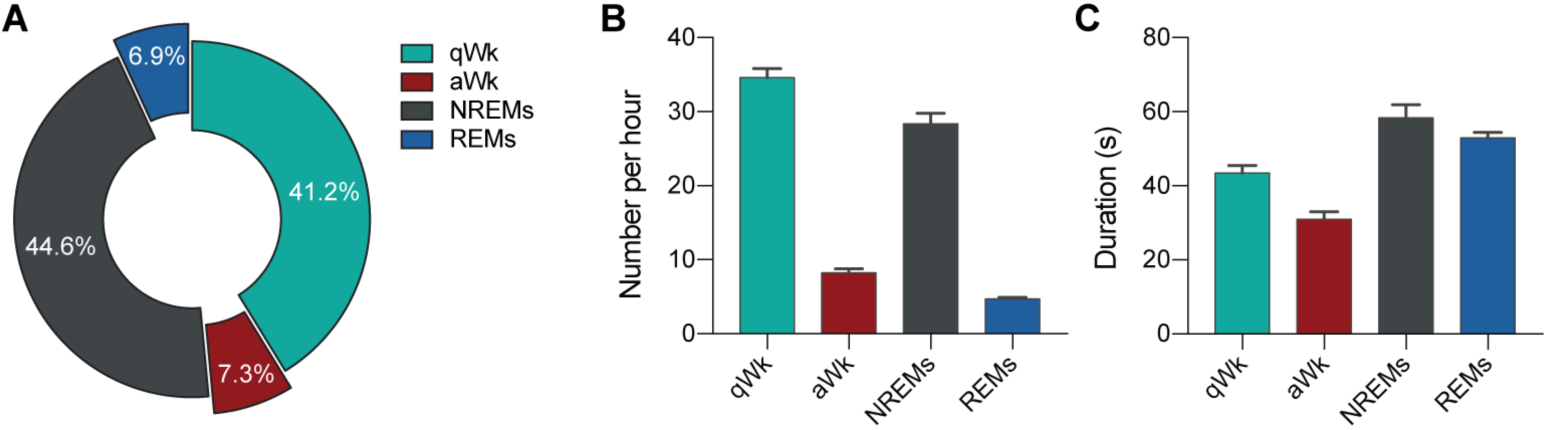
Quantification of vigilance states obtained from 11 animals, 24 h each. (*A*) Percentage of time at each state. On average animals spent almost equal time periods in aWk and REMs. (*B*) Number of episodes per hour for each state. (C) Duration of episodes for each state. Bars represent mean ±SEM.

**Table S1.** Theta-MG coupling and inter-coupling interval characteristics during aWk and REMs for CA3, CA1, subiculum, and PCx recordings. The reported values are mean ±SEM from 11 mice (REMs vs. aWk: **p < 0.01, ***p < 0.001, Wilcoxon test).

|  | CA3 |  | CA1 |  | Sub |  | PCx |  |
| --- | --- | --- | --- | --- | --- | --- | --- | --- |
|  | aWk | REMs | aWk | REMs | aWk | REMs | aWk | REMs |
| Theta-MG Coup. ($\times 10^{-3}$) | 5.4 ± 0.8 | 7.8 ± 1.3 | 8.3 ± 1.2 | 12 ± 2.3 | 7.1 ± 0.7 | 15 ± 1.8 ** | 13 ± 1.2 | 25 ± 2.3 *** |
| Inter-coupling interval (s) | 15.9 ± 0.4 | 16.9 ± 0.4 | 15.2 ± 0.4 | 17.7 ± 0.4 | 15.8 ± 0.3 | 17.8 ± 0.4 | 15.8 ± 0.4 | 16.9 ± 0.4 |

**Table S2.** Analyses from pairwise combination of recording sites. Overlap ratio and correlation correspond to theta-MG coupling events of recording sites during aWk and REMs (mean ±SEM; n = 11 animals; REMs vs. aWk: *p < 0.05, **p < 0.01, ***p < 0.001, Wilcoxon test). Theta phase coherence reports coherency values during baseline and optogenetic silencing of MS^GABA^ cells (mean ±SEM; n = 5 animals, 10 trials each; silencing vs. baseline: ***p < 0.001, Wilcoxon test).

|  | Overlap ratio |  | Correlation |  | Theta phase coherence |  |
| --- | --- | --- | --- | --- | --- | --- |
|  | aWk | REMs | aWk | REMs | Baseline | Silencing |
| CA3-CA1 | 0.13 ± 0.01 | 0.17 ± 0.02 ** | 0.09 ± 0.02 | 0.19 ± 0.04 ** | 0.76 ± 0.03 | 0.44 ± 0.04 *** |
| CA3-Sub | 0.12 ± 0.01 | 0.19 ± 0.01 ** | 0.09 ± 0.02 | 0.27 ± 0.05 ** | 0.81 ± 0.04 | 0.48 ± 0.05 *** |
| CA3-PCx | 0.14 ± 0.01 | 0.16 ± 0.02 | 0.12 ± 0.03 | 0.26 ± 0.06 * | 0.70 ± 0.03 | 0.30 ± 0.03 *** |
| CA1-Sub | 0.19 ± 0.02 | 0.30 ± 0.02 ** | 0.21 ± 0.03 | 0.44 ± 0.04 ** | 0.88 ± 0.02 | 0.64 ± 0.04 *** |
| CA1-PCx | 0.14 ± 0.01 | 0.21 ± 0.02 ** | 0.12 ± 0.03 | 0.32 ± 0.05 *** | 0.78 ± 0.04 | 0.44 ± 0.04 *** |
| Sub-PCx | 0.19 ± 0.02 | 0.25 ± 0.02 * | 0.24 ± 0.05 | 0.42 ± 0.04 ** | 0.83 ± 0.03 | 0.51 ± 0.04 *** |

**Table S3.** Theta-MG coupling, theta power, and MG power during middle and late segments of REMs. The reported values were normalized and compared with early segment of REMs (mean ±SEM; n = 11 animals; **p < 0.01, ***p < 0.001, Wilcoxon test).

|  | CA3 |  | CA1 |  | Sub |  | PCx |  |
| --- | --- | --- | --- | --- | --- | --- | --- | --- |
|  | Middle REMs | Late REMs | Middle REMs | Late REMs | Middle REMs | Late REMs | Middle REMs | Late REMs |
| Theta-MG Coup. ($\times 10^{-3}$) | 1.36 ± 0.05 *** | 1.58 ± 0.06 *** | 1.27 ± 0.04 *** | 1.39 ± 0.04 *** | 1.17 ± 0.04 *** | 1.24 ± 0.04 *** | 1.21 ± 0.03 *** | 1.29 ± 0.03 *** |
| Theta power | 0.95 ± 0.02 | 0.97 ± 0.02 | 1.03 ± 0.02 | 1.03 ± 0.02 | 1.06 ± 0.02 | 1.11 ± 0.02 ** | 1.04 ± 0.01 | 1.04 ± 0.01 |
| MG power | 1.18 ± 0.01 *** | 1.23 ± 0.02 *** | 1.21 ± 0.01 *** | 1.26 ± 0.02 *** | 1.2 ± 0.01 *** | 1.24 ± 0.02 *** | 1.29 ± 0.01 *** | 1.37 ± 0.02 *** |

**Table S4.** Analyses of theta-gamma coupling characteristics during optogenetic silencing of MS^GABA^ cells within REMs. All reported values were normalized to baseline (mean ±SEM; n = 5 animals, 10 trials each; silencing vs. baseline: ***p < 0.001, Wilcoxon test).

|  | Theta power | MG power | HG power | UG power | Theta-MG Coup. | Theta-HG Coup. | Theta-UG Coup. |
| --- | --- | --- | --- | --- | --- | --- | --- |
| CA3 | 0.53 ± 0.04 *** | 0.91 ± 0.02 | 0.92 ± 0.03 | 0.92 ± 0.02 | 0.29 ± 0.11 *** | - | 0.61 ± 0.07 *** |
| CA1 | 0.50 ± 0.05 *** | 0.91 ± 0.04 | 0.90 ± 0.03 | - | 0.21 ± 0.04 *** | - | - |
| Sub | 0.55 ± 0.08 *** | 0.92 ± 0.05 | 0.95 ± 0.04 | 0.93 ± 0.02 | 0.24 ± 0.09 *** | - | 0.2 ± 0.04 *** |
| PCx | 0.53 ± 0.02 *** | 0.94 ± 0.04 | 0.93 ± 0.03 | - | 0.27 ± 0.09 *** | 0.27 ± 0.08 *** | - |

